# Forest growth responds more to air pollution than soil acidification

**DOI:** 10.1101/2021.08.23.457413

**Authors:** Jakub Hruška, Filip Oulehle, Tomáš Chuman, Tomáš Kolář, Michal Rybníček, William H. McDowell

**Affiliations:** Global Change Research Institute, Czech Academy of Sciences, Bělidla 986/4a, 603 00 Brno, Czech Republic; Czech Geological Survey, Klárov 3, 118 21, Praha 1, Czech Republic; Department of Wood Science and Technology, Faculty of Forestry and Wood Technology, Mendel University in Brno, Zemědělská 3, Brno, 613 00, Czech Republic; Department of Natural Resources and the Environment, University of New Hampshire, Durham, 03824, NH, United States

**Author notes:** These authors contributed equally to this work (Jakub Hruška, Filip Oulehle, Tomáš Kolář). These authors also contributed equally to this work (Tomáš Chuman, Michal Rybníček, William H. McDowell).

## Abstract

The forests of central Europe have undergone remarkable transitions in the past 40 years as air quality has improved dramatically. Retrospective analysis of Norway spruce (*Picea abies*) tree rings in the Czech Republic shows that air pollution (e.g. SO_2_ concentrations, high acidic deposition to the forest canopy) plays a dominant role in driving forest health. Extensive soil acidification occurred in the highly polluted “Black Triangle” in Central Europe, and upper mineral soils are still acidified. In contrast, acidic atmospheric deposition declined by 80% and atmospheric SO_2_ concentration by 90% between the late 1980s and 2010s. Annual tree ring width (TRW) declined in the 1970s and subsequently recovered in the 1990s, tracking SO_2_ concentrations closely. Furthermore, recovery of TRW was similar in unlimed and limed stands. Despite large increases in soil base saturation, as well as soil pH, as a result of repeated liming starting in 1981, TRW growth was similar in limed and unlimed plots. TRW recovery was interrupted in 1996 when highly acidic rim (originating from more pronounced decline of alkaline dust than SO_2_ from local power plants) injured the spruce canopy, but recovered soon to the pre-episode growth. Across the long-term site history, changes in soil chemistry (pH, base saturation, Bc/Al soil solution ratio) cannot explain observed changes in TRW at the two study sites at which we tracked soil chemistry. Instead, statistically significant recovery in TRW is linked to the trajectory of annual SO_2_ concentrations or sulfur deposition at all three stands.

## 1. Introduction

Central Europe was heavily polluted by SO_2_ originating from burning high sulfur lignite for electricity generation [1, 2]. In the so-called “Black Triangle” region on the border of Germany (former DDR), Poland and the Czech Republic (former Czechoslovakia), massive forest dieback occurred starting in the 1960s with a peak in the 1970s and 1980s. In the Czech Republic alone, 1.5 million ha of forest were heavily damaged and about 40 000 ha of mainly Norway spruce (*Picea abies*) stands died or were salvaged logged due to air pollution [3], which represents ca. 9.5 million cubic meters of wood [2]. The decline in SO_2_ emission during the 1990s was one of the great “success stories” in environmental protection worldwide [4, 5]. Czech SO_2_ emissions declined from 3 150 Mkg of SO_2_ to 86.6 Mkg in 2017 [2], representing a decline of more than 97%. Annual ambient SO_2_ concentration in the Czech part of the Black Triangle has declined from up to 130 μg·m^-3^ measured in the 1970s to less than 10 μg·m^-3^ at present [2]. The estimated median total deposition of S in the current Czech Republic peaked in 1979 (41 kg S ha^−1^·yr^−1^) and then declined to 7.3 kg S ha^−1^·yr^−1^ in 2012 [6]. Recent estimates show that S deposition had fallen even further by 2017 (5.4 kg S ha^−1^·yr^−1^).

Acid deposition results in elevated inorganic Al and H^+^ in soil solution, especially in soils with low base saturation (<20%). For forests, where the toxicity of aluminium to tree roots is considered to be critical, the Al^3+^ to Ca^2+^ ratio in soil water has become the main driver of damage to roots [7, 8] or crown defoliation and transformation. High inorganic Al in soil solution can also impact tree fine root growth and functioning [9].

Foliar injury by air pollutants has also been identified as a reason for forest dieback in highly polluted areas such as the Black Triangle [10], and high concentrations of SO_2_ and ozone result in canopy decline [11]. Deterioration of cuticular waxes and leaching of nutrients from the canopy [12] leading to chlorosis and decline of photosynthesis is thought to drive the canopy decline and contribute to forest dieback. Widespread and quick declines of Norway spruce in the Black Triangle in the 1970s were usually also connected with climatic episodes when highly acidic fog or rim during winter inversions seriously damaged spruce canopy [1].

One of the best opportunities for evaluating the future capacity of individual species is a retrospective analysis of past growth responses to climate and pollution [13, 14]. High-resolution long-term proxies, where annual changes can be delineated, are needed for describing previous environmental variability [15]. Annual rings produced by temperate tree species provide a retrospective record of tree growth in response to past environmental conditions [16, 17, 18].

In this paper, we examined the growth response of Norway spruce (*Picea abies*) to (i) long-term air pollution by SO_2_ (ii); acidic deposition; and (iii) soil chemistry in the formerly extremely SO_2_-polluted Ore Mts. in the northern Czech Republic, to determine the most important factors affecting forest vitality during the past 50 years.

## 2. Site description

The study sites are spruce stands located in the Ore Mountains (Krušné hory in Czech, Erzgebirge in German), northwestern Czech Republic (Fig. 1). All Norway spruce (*Picea abies*) stands are at a similar altitude (Tab. 1) and are exposed to similar climatic conditions. The average annual temperature in the region is 7.1°C, and the average annual precipitation is 1110 mm (2005–2017).

**Figure 1.**
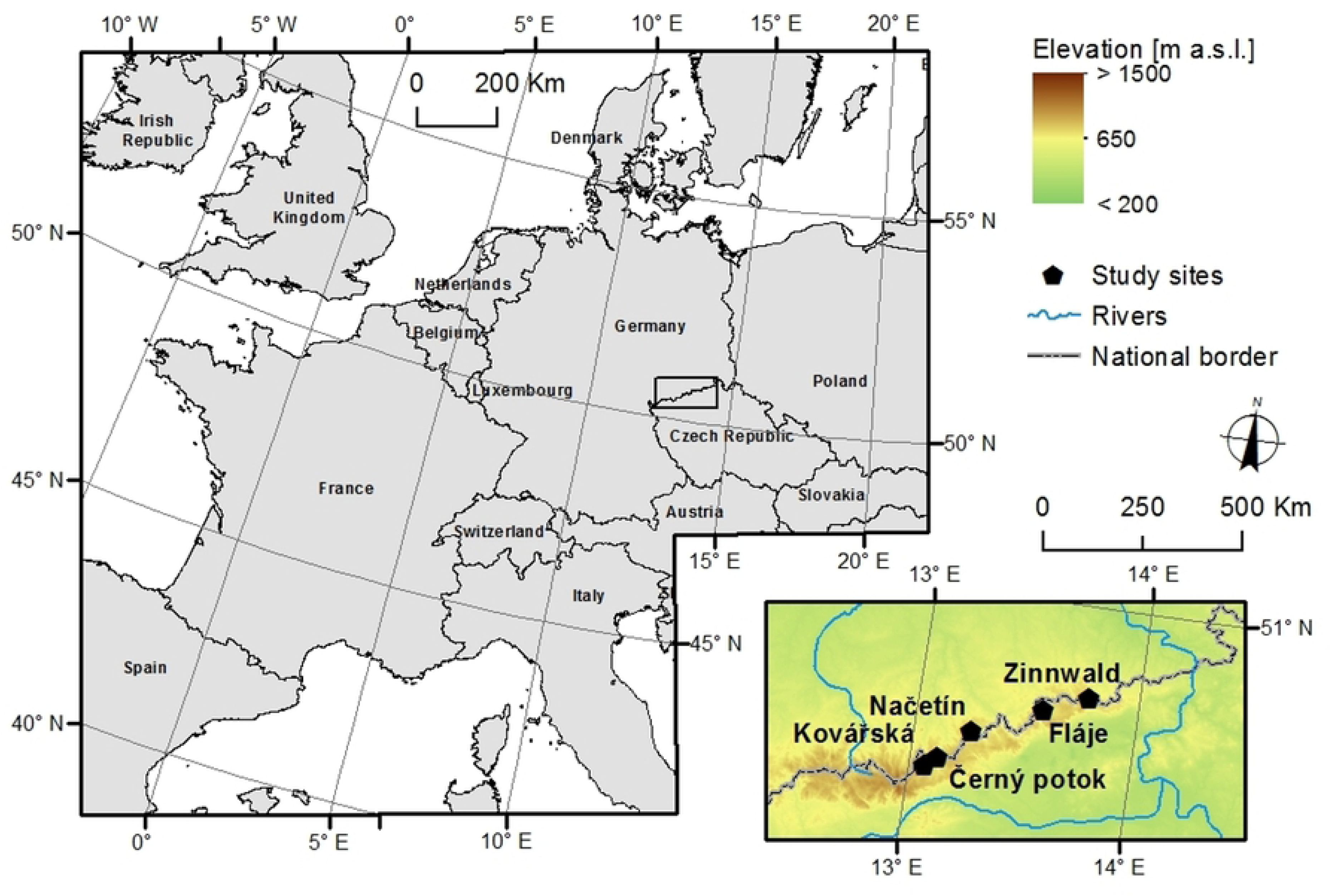
Study site locations in Ore Mts., Czech Republic.

**Table I.** Characteristics of studied plots in Ore Mts., Czech Republic.

| Site | Coordinates | Bedrock | Soil | Altitude<br>(m a.s.l.) | Forest age<br>(years)* | Liming (year,<br>dose in tons.ha<sup>-1</sup>) |
| --- | --- | --- | --- | --- | --- | --- |
| Načetín | 50° 33' 23''<br>13° 15' 12'' | Gneiss | Dystric<br>cambisol | 775 | 95 | 2007**, 3.0 |
| Kovářská | 50° 29' 53''<br>13° 07' 20'' | Gneiss | Dystric<br>cambisol | 786 | 116 | 1981, 1.5<br>1988, 2.0<br>1997, 3.0 |
| Černý<br>potok | 50° 28' 09''<br>13° 03' 54'' | Gneiss | Dystric<br>cambisol | 768 | 69 | 1981, 1.5<br>1988, 2.0<br>1997, 3.0 |
| Fláje | 50° 40' 48''<br>13° 34' 12'' | Basalt | Ranker | 860 | 69 | 1981, 1.5<br>1988, 2.0<br>1997, 3.0 |

\*first identified tree ring +10 years
\*\*Načetín limed only

The Načetín experimental forest stand has been studied since the late 1980s as part of research into the phenomenon of acid rain [19, 20, 21]. Experimental liming by 3 tons per hectare of dolomitic limestone occurred in an experimental plot of Načetín site in 2007, thus “Načetín control” and “Načetín liming” will be used to identify different subplots.

Tree rings were sampled at each plot, although soil chemistry was not analysed at Černý potok as the site position, bedrock, and soils were almost identical with the nearby Kovářská site.

## 3. Methods

### 3.1. Tree-ring sampling and chronology development

We randomly sampled 15 spruce trees at each forest site. Considering the lack of direction-specific effects on variability in tree radial growth [22], one core per tree was extracted using a Pressler borer (Haglof Company Group) at breast height (1.3 m). To avoid compression wood, the cores were sampled in a direction parallel to the slope. All samples were measured using a VIAS TimeTable device with a measuring length of 78 cm and resolution <1/100 mm (©SCIEM, Vienna, Austria). The TRW series were measured (accuracy of 0.01 mm) and cross-dated using PAST4 [23]. Missing and false rings were corrected using both PAST4 [23] and COFECHA [24].

To remove non-climatic, age-related growth trends from the raw tree ring width (TRW) series as well as other non-climatic factors (e.g., competition), we applied cubic smoothing splines with a 50% frequency cutoff at 100 years [25] using ARSTAN software [26]. We used this method to preserve interannual to multi-decadal growth variations [27]. TRW indices were calculated as residuals after applying an adaptive power transformation to the raw measurement series [28]. The site chronologies were calculated using bi-weight robust means. Similarities among the site TRW chronologies were assessed using statistical criteria (correlation coefficient; Gleichläufigkeit [29]) and visual comparison. Negative extremes were defined by years in which the standardised TRW chronology (period replicated >20 TRWi series) exceeded the -1.0 multiple of a standard deviation (SD).

### 3.2. Soil analyses

Soils were sampled in 2018 by excavating four 0.5 m^2^ quantitative pits using the method described by Huntington et al. [30]. The O_l_ (litter) and O_f_+O_h_ (fermented + humified) horizons were sampled together. Mineral soil was collected for the depths of 0–10, 10–20 and 20–40 cm. The soil samples were weighed and then sieved after air-drying (mesh size of 5 mm for organic horizons and 2 mm for mineral horizons). Soil moisture was determined gravimetrically by drying at 105^◦^C. Soil pH was measured using a glass combination electrode in a water extract. Exchangeable cations were analysed in 0.1 M BaCl_2_ extracts by the flame atomic absorption spectrometry (FAAS). Cation exchange capacity (CEC) was calculated as the sum of exchangeable Ca, Mg, Na, K and total exchangeable acidity (TEA). Base saturation (BS) was determined as the fraction of CEC associated with base cations. Total carbon (C) and total nitrogen were determined using an elemental analyser. Soil water chemistry at Načetín was sampled monthly since 1994 using quartz and Teflon suction lysimeters (Prenart) from the depth of -30 cm of mineral soil.

### 3.3. MAGIC model

MAGIC (Model of Acidification of Groundwater in Catchments) was designed to reconstruct past and predict future drainage water and soil chemistry [31, 32, 33]. MAGIC is a lumped-parameter model of intermediate complexity which calculates for annual time steps the concentration of major ions under the assumption of simultaneous reactions involving SO_4_ adsorption, cation exchange, dissolution-precipitation-speciation of aluminium and dissolution-speciation of inorganic and organic C. MAGIC accounts for the mass balance of major ions in the soil by bookkeeping the fluxes from atmospheric inputs, chemical weathering, net uptake in biomass and loss to runoff. Water fluxes, wet and dry atmospheric deposition, net vegetation uptake, weathering, and a description of organic acids are required as external inputs to MAGIC. For calibration procedures, see S1 Annex.

## 4. Results and Discussion

### 4.1. Atmospheric chemistry and deposition

Historically, sulfur emission and deposition were associated with mining and burning brown coal (lignite) from the nearby Bohemian coal basin (Fig. 2) since the second half of the 19^th^ century. The period after World War II was accompanied by a massive production of energy by burning this high S content lignite in the local power plants. Coal mining, as well as SO_2_ emissions (Fig. 3), peaked in the 1980s. The decrease since then can be attributed in part to the declining volume of coal mined after 1989, when the extensive industrial activity in former Czechoslovakia declined with a change in political structure. Coal mining declined from its peak in 1982 (130 Mt) to 58 Mt in 2017 [2].

**Figure 2.**
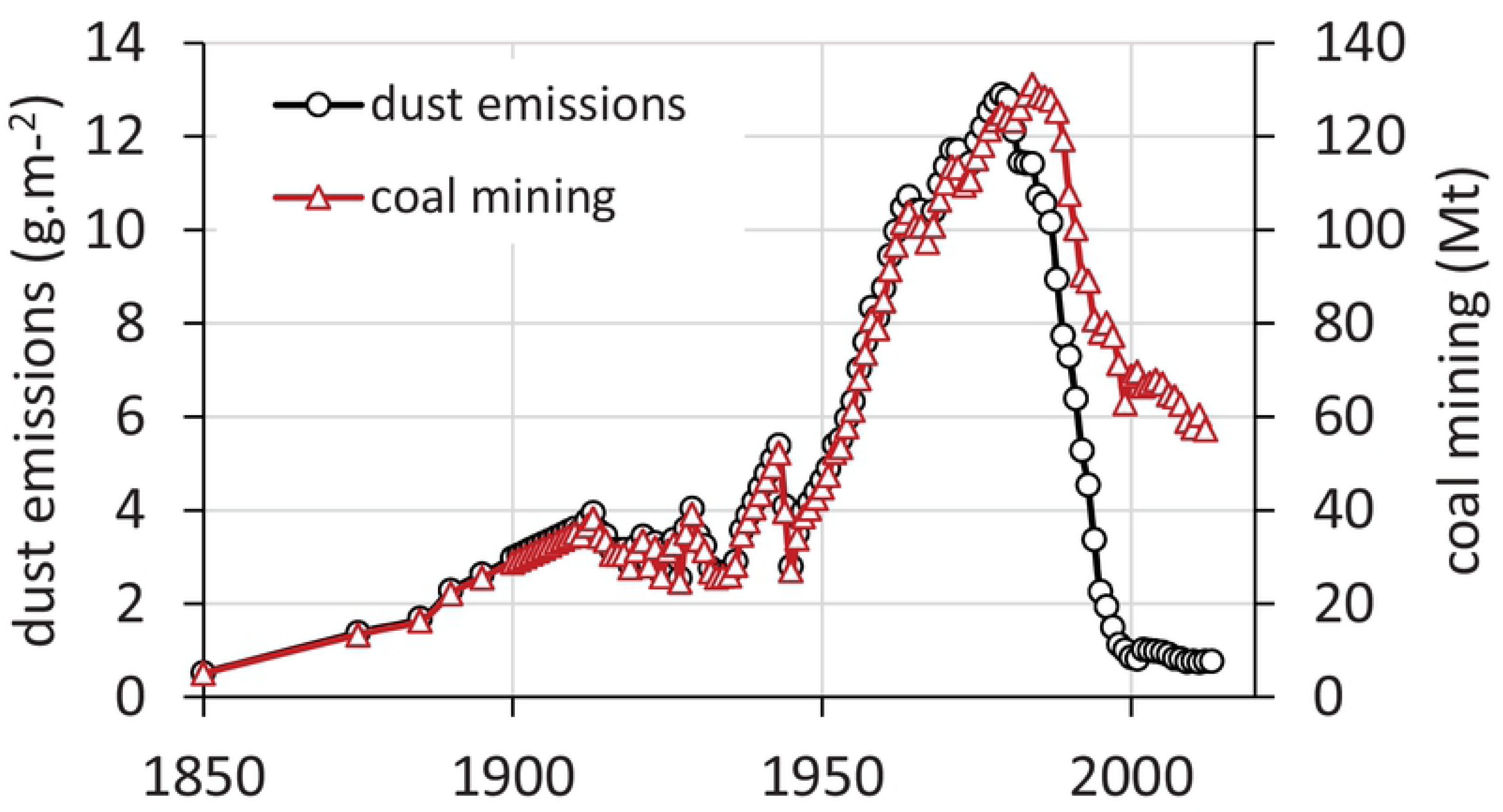
Coal mining and dust emissions in the Czech Republic between 1850-2017.

**Figure 3.**
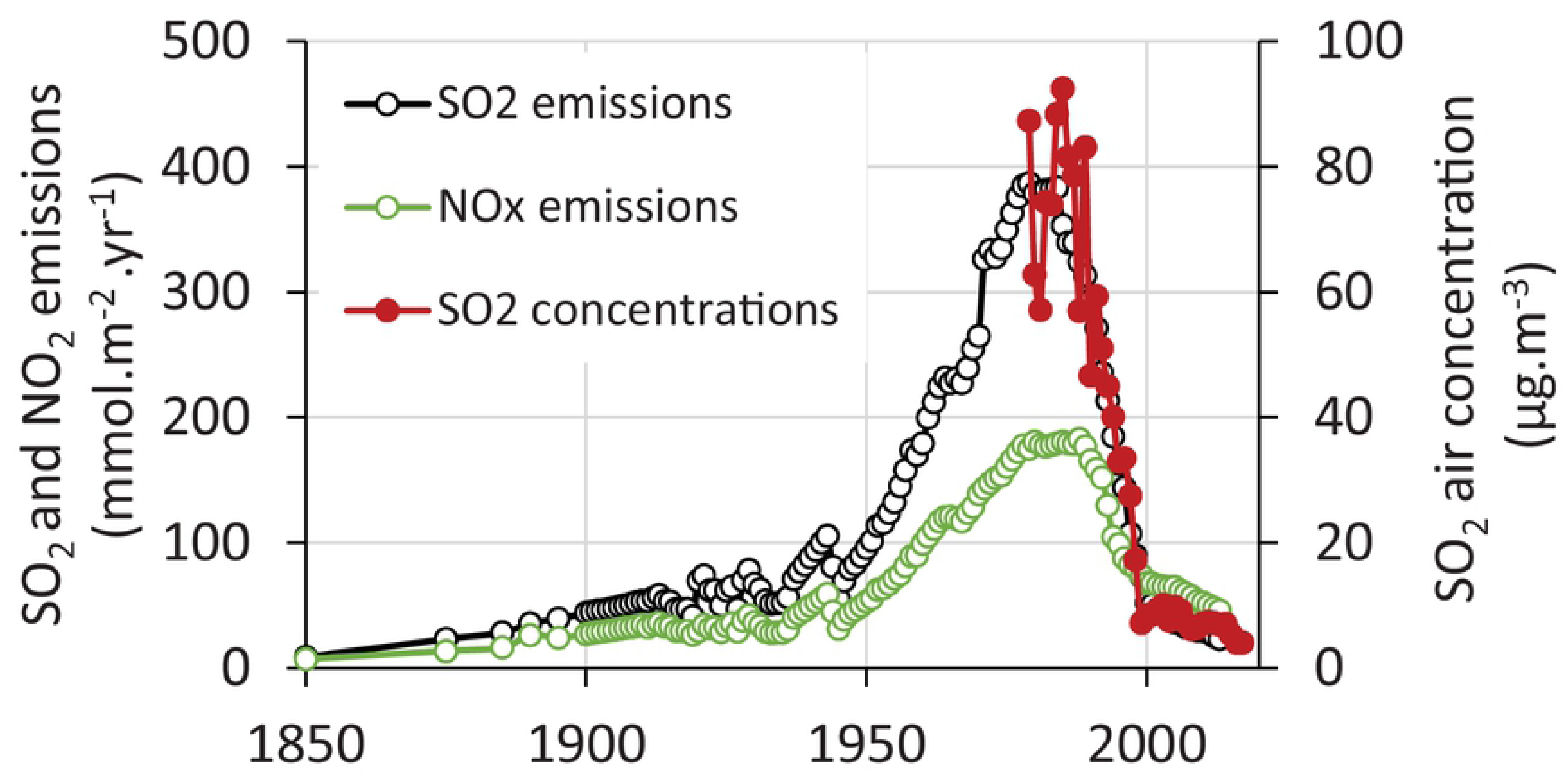
SO_2_ emissions (Czech Republic) between 1850-2017 and ambient annual SO_2_ concentrations measured at Cínovec (Zinwald) at the Czech/German border between 1979-2017.

There was a distinct break in SO_2_ emissions in 1993 when the first power plants in the Czech Republic were equipped with flu-gas desulfurisation (Fig. 3). This process was completed in 1999. As a result, SO_2_ emissions declined from a peak of 395 mmol·m^-2^·yr^-2^ in 1982 to 25 mmol·m^-2^·yr^-2^ in 1999. Since then, it fell to 10 mmol·m^-2^·yr^-2^ in 2017 (Fig. 3) due to the continuous decline in coal mining (Fig. 2). Modelled S emissions coincide very well with measured ambient SO_2_ concentrations. The longest record in the region is available from Zinwald station [34] on the Czech-German border on the ridge of the Ore Mts. (Fig. 1). Annual concentrations peaked in the mid-1980s at 93 μg·m^-3^ (Fig. 3). As SO_2_ emissions declined, ambient SO_2_ concentrations declined proportionally to ca. 10 μg·m^-3^ in the late 1990s and 4 μg·m^-3^ in 2017.

Dust emissions (Fig. 2) increased similarly to coal mining activities [6]. Dust was rich in base cations (primarily Ca) and partly neutralised precipitation acidity (Fig. 4). Despite this neutralisation, precipitation pH was about 4.2 during the 1970s and 1980s [35] (Fig. 4) and has risen slowly since the 1990s to values >5.0 in recent years. Electrostatic removal of dust from power plants started in the 1980s when it peaked at around 12 g·m^-2^. It declined sharply to <1 g·m^-2^ until the late 1990s (Fig. 2). As dust removal was effective earlier than SO_2_ removal (Fig. 3), the ratio of SO_2_/dust in the atmosphere peaked in the 1990s (Fig. 4). This peak in SO_2_/dust was accompanied by highly acidic episodes like the formation of acidic rime ice in the winter of 1995/1996. The acidity of the rime developed on the spruce canopy at Načetín was measured to be pH<3 (lowest pH=2.33 was recorded in January/February 1996). Ambient SO_2_ concentration was 2 300 μg·m^-3^ at Rudolice, 20 km east of Načetín (February 2^nd^ 1996) as a result of repeated inversions occurring on the ridge of the Ore Mts. [36].

**Figure 4.**
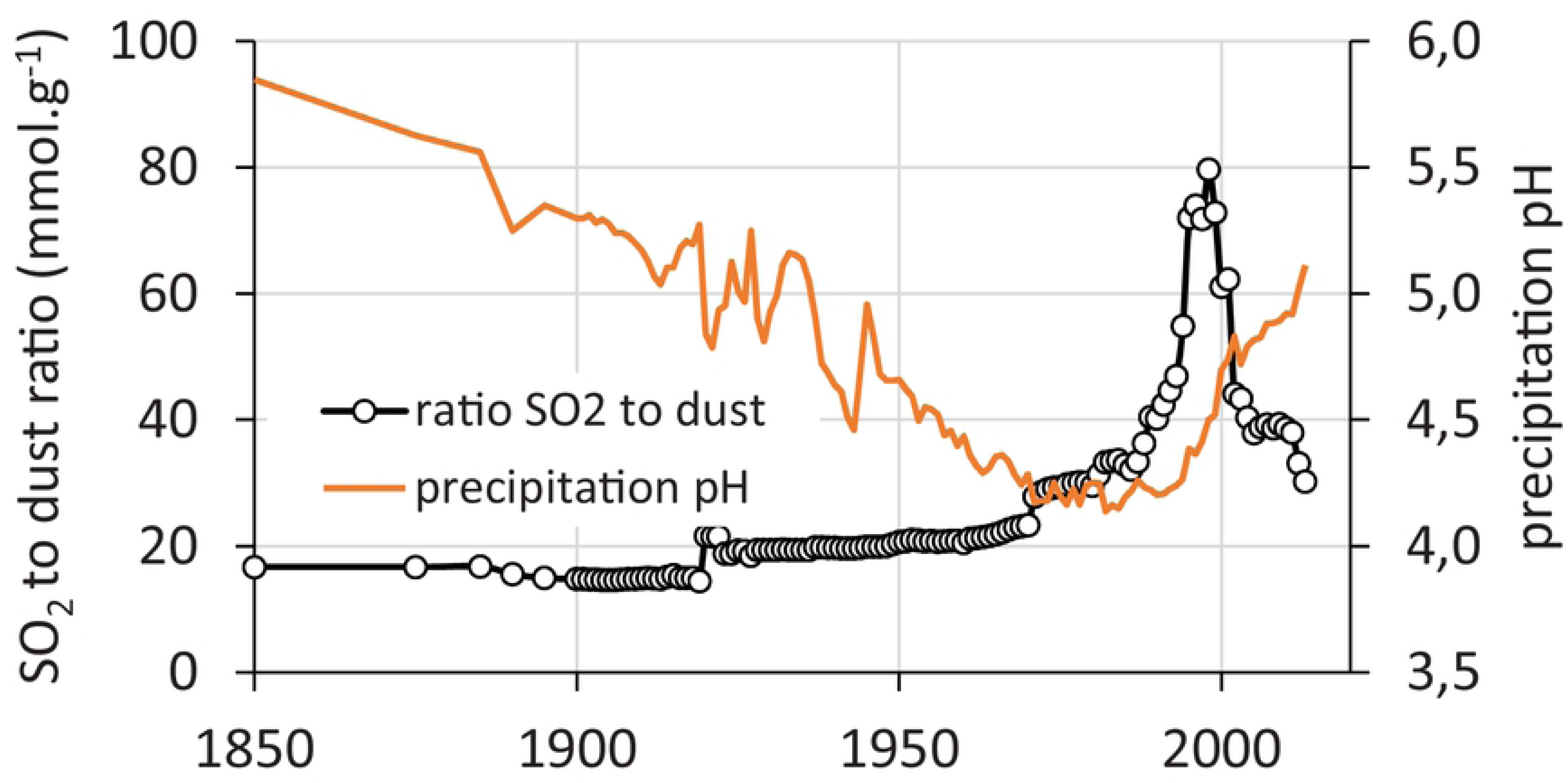
Modelled and measured annual precipitation pH and SO_2_/dust ratio (1850-2017) for the Czech Republic. Modelled values from 1850 to 2013 for pH, 1850 to 2013 for SO_2_/dust ratio.

Based on the coal mining and measured sulfur deposition data between the 1990s and 2010s from the Czech Republic, Germany and Poland [6], constructed a simple statistical model for reconstructing and predicting historic S deposition. This model estimated the deposition of S in throughfall for sites in the Ore Mts (Fig. 5) for the period 1850-2017. All sites showed synchronous patterns, with the highest deposition estimated for Fláje (ca. 550 meq·m^-2^·yr^-1^ in the 1980s), but other sites (Kovářská and Načetín) also received very high loads (ca. 480 meq·m^-2^·yr^-1^). Present deposition (60-70 meq·m^-2^·yr^-1^) is equal to the historical values estimated for the second half of the 19^th^ century and representing an 80-90% decline (Fig. 5).

**Figure 5.**
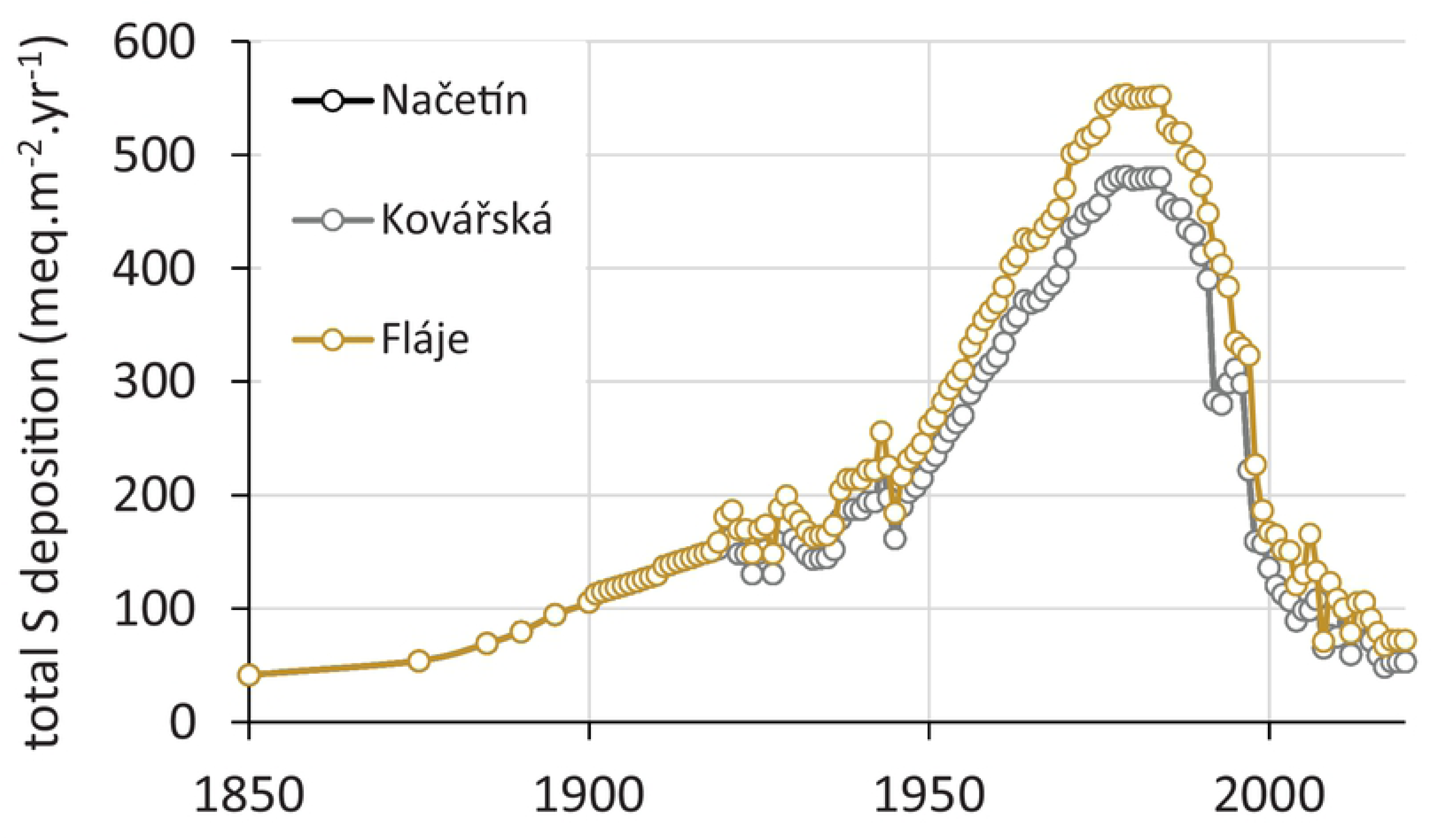
Modelled (1850-1993) and measured (1994-2017) annual sulfur deposition at study sites in the Ore Mts.

### 4.2. Soil chemistry

#### 4.2.1. Long-term changes at Načetín

Since 1994, four soil sampling campaigns were undertaken at the Načetín control site. Despite a significant decrease in atmospheric deposition of sulfur (Fig. 5), soil chemistry exhibited only limited changes. Soil base saturation (BS) has increased only in the organic layer between 1994 and 2003 and has stayed at a similar level of 33-37% since 2003 (Fig. 6). Mineral soil BS did not change significantly between 1994 and 2008, but declined in 2018 to the lowest measured levels, with values of only 3% of saturation at a depth of 20-40 cm compared to the initial value of 6% at that depth in 1994 (Fig. 6). Total exchangeable acidity (TEA) declined significantly in the humus horizon (from 125 mmol_c_·kg^-1^ in 1994 to 67 mmol_c_·kg^-1^ in 2018). It decreased markedly in the upper mineral soil (0-10 cm) but did not change in deeper mineral soil (Fig. 6). Soil pH did not exhibit consistent trends over time and increased from 3.6 in the forest floor up to 4.4 in the B horizon (20-40 cm). Exchangeable calcium has declined similarly to TEA, with significant declines in the humus horizon (from 600-700 mg·kg^-1^ in 1994 to about 450 in 2018) and in the upper mineral soil (from 70 to 15 mg·kg^-1^). Similar sharp declines were observed in deeper mineral horizons, with exchangeable Ca dropping from about 30 to 7 mg·kg^-1^ at 10-20 cm and from 20 to 4 mg·kg^-1^ at 20-30 cm (Fig. 6). Patterns for exchangeable Mg were similar to those of Ca (data not shown).

**Figure 6.**
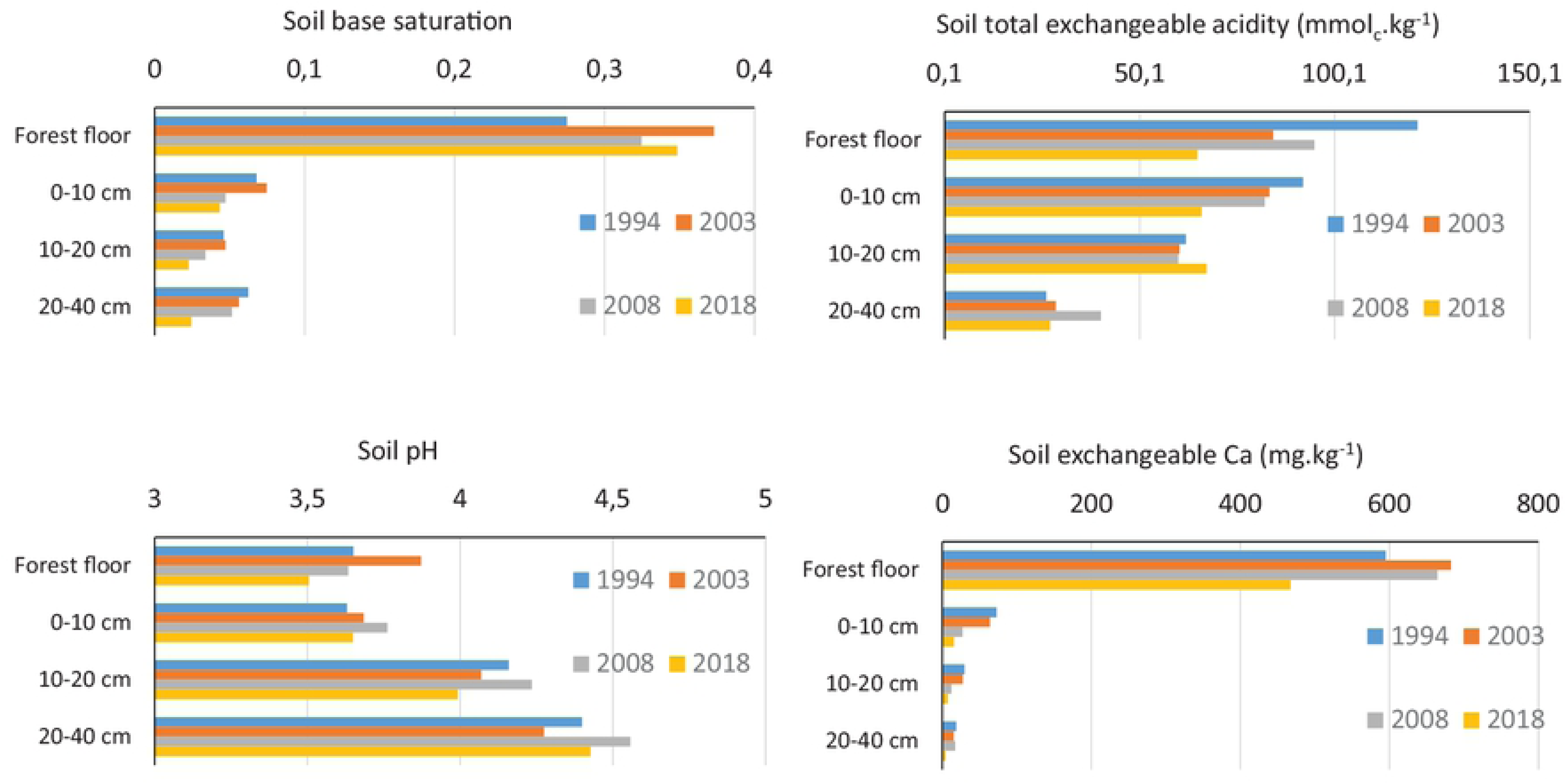
Long-term changes of soil chemistry at the Načetín control research site (1994-2018).

#### 4.2.2. Variation in soil chemistry among sites

Among our study sites, the most acidic was the Načetín control plot (exchangeable pH = 3.4 at FH horizon) followed by Načetín limed plot, Kovářská and Fláje (4.7 at FH horizon, Fig. 7). Soil pH showed the typical pattern being most acidic in the organic FH horizons and the highest exchangeable pH was observed in the deeper mineral soil. Soil base saturation has declined at all sites from L horizon down to mineral 20-40 cm. The lowest BS was observed for the Načetín control plot (70% in L horizon and 3% in 20-40 cm) and highest at Fláje (95% in L and 39% in 20-40 cm respectively).

**Figure 7.**
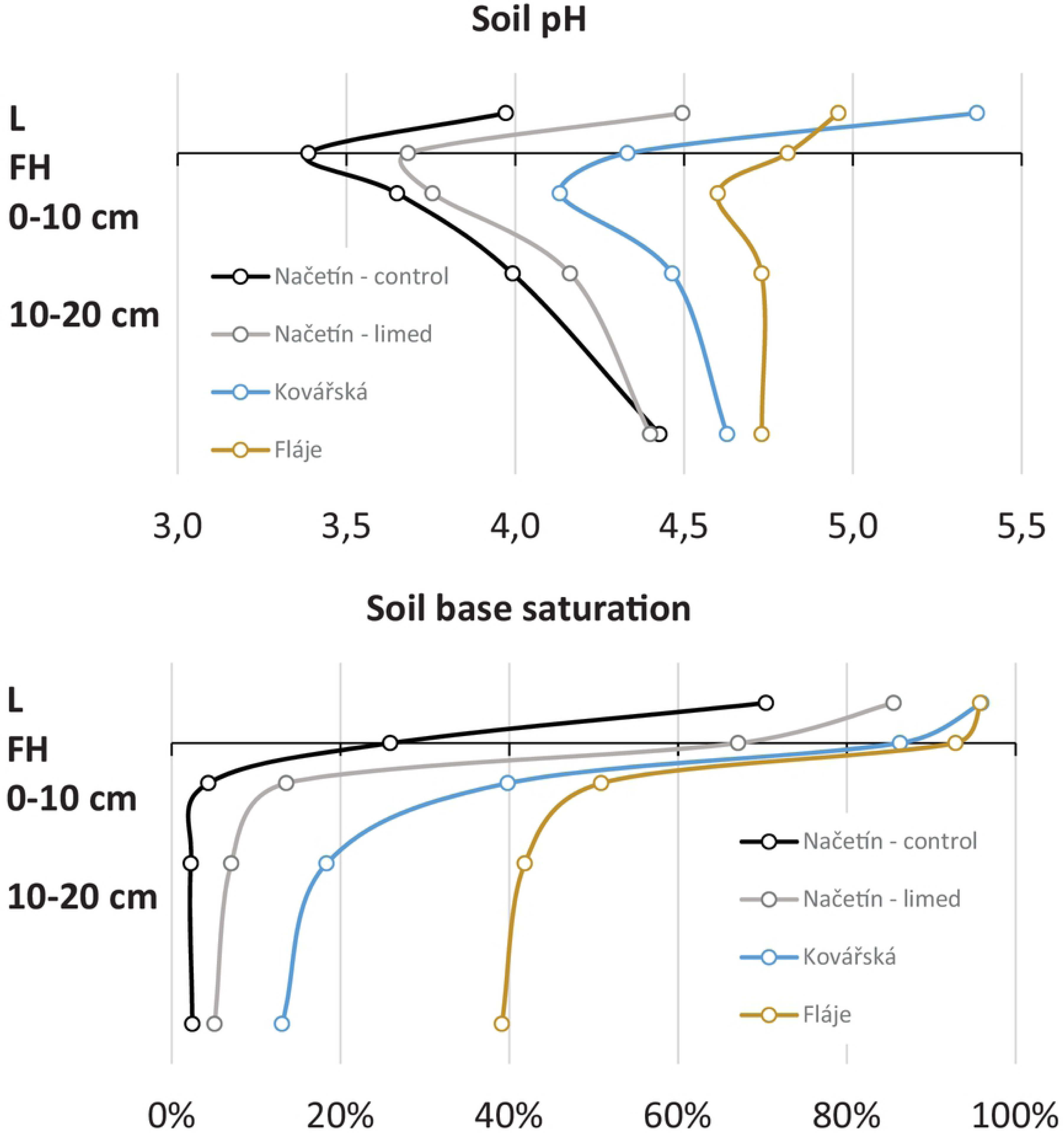
Soil pH and base saturation in 2018.

Soil base saturation follows liming history as well as geological settings. The second most acidic site (Načetín limed, underlined by gneiss) was experimentally limed in 2007 (Table I). Other sites Kovářská (gneiss) and Fláje (basalt) were limed three times (Table I) between 1981 and 1997 [37, 38] by a cumulative dose of 6.5 t.ha^-1^. As a result, soil pools of Ca were 8x higher, and the Mg pool was 11x enriched (Table 2) at Kovářská, even though it is underlain by similar bedrock (gneiss) as Načetín. Similar levels of enrichment were observed at Fláje, which is underlain by basalt. As Načetín was limed in 2007 by a lower dose (3 t·ha^-1^), Ca enrichment was only 3x and Mg 4x (Tab. 2), reflecting slow kinetics of the limestone dissolution (S1 Annex).

**Table II.**
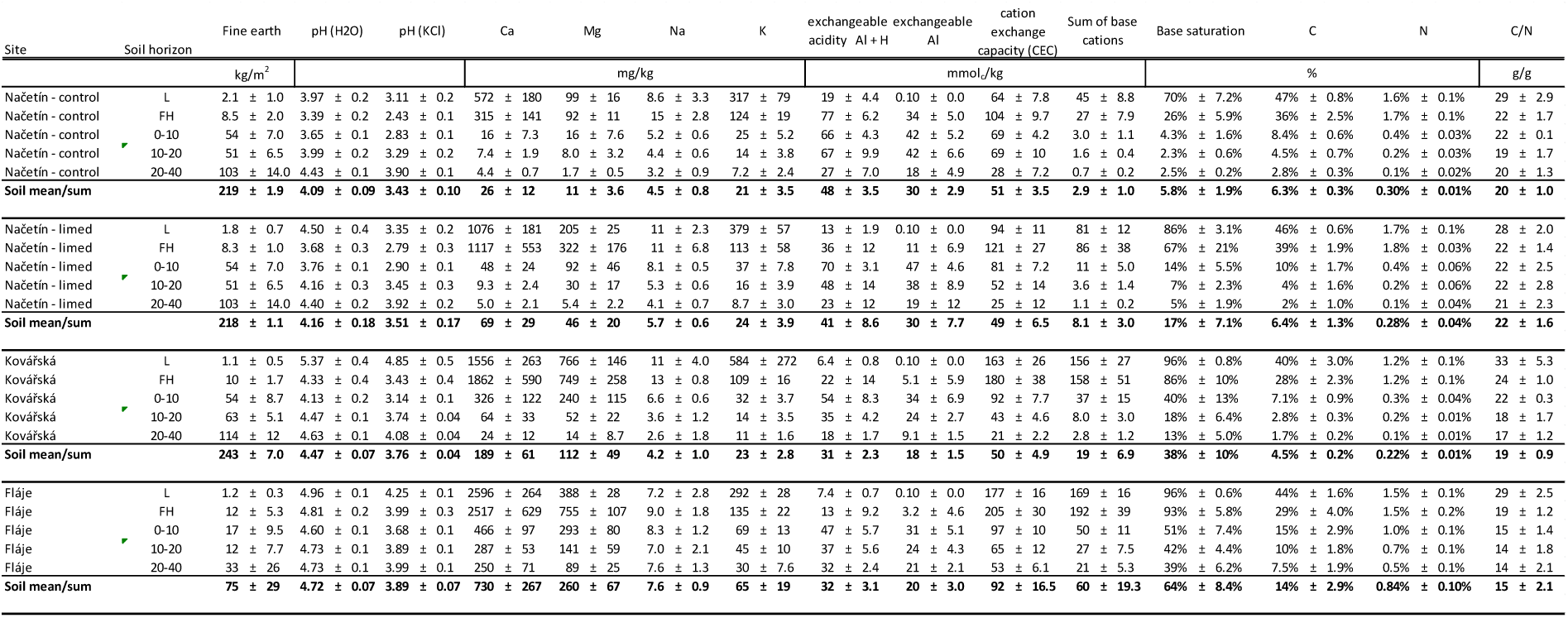
Pools of base cations, carbon and nitrogen measured in 2018.

#### 4.2.3. Estimate of historical soil chemistry development

As tree rings reflected forest damage and the physiological stress in the past, retrospective soil chemistry development was needed for disentangling the effects of ambient air chemistry, atmospheric deposition and soil chemistry on tree growth. According to the MAGIC model estimates (S1 Annex), soil base saturation declined during the 19^th^ and most of the 20^th^ century due to increasing acidic deposition (Fig. 5) at all of our study sites (Fig. 8), indicating an ongoing depletion of the soil pool of exchangeable base cations. At Načetín control, where liming was not applied, the MAGIC simulations showed significant depletion, from an estimated base saturation of 19% in 1850 to 6% in 2018. The projected recovery, with acid deposition assumed to be unchanged from levels in 2018, revealed only minor improvements of soil base saturation (7% in 2050). This indicates a balance between base cations inputs (weathering + deposition) and outputs (uptake + leaching) for future decades. Other limed sites exhibited similar patterns until liming when base saturation increased significantly (Fig. 8). Načetín limed revealed now BS of 16.5%, not far from the preindustrial estimate (19%). Future scenarios predict base saturation of 27% in 2050. A more pronounced liming effect was observed and modelled for Kovářská where the preindustrial estimate was 19% but measured BS was 38.5% in 2018. This high base saturation was due to earlier liming (beginning in 1981) and a higher cumulative dose (Table I). More alkaline bedrock with a high weathering rate (see S1 Annex) at Fláje resulted in preindustrial base saturation of 45% (Fig. 8). Acid deposition lowered BS to 15% in 1981 and subsequent liming raised BS to 64% in 2018, significantly higher than the preindustrial estimate.

**Figure 8.**
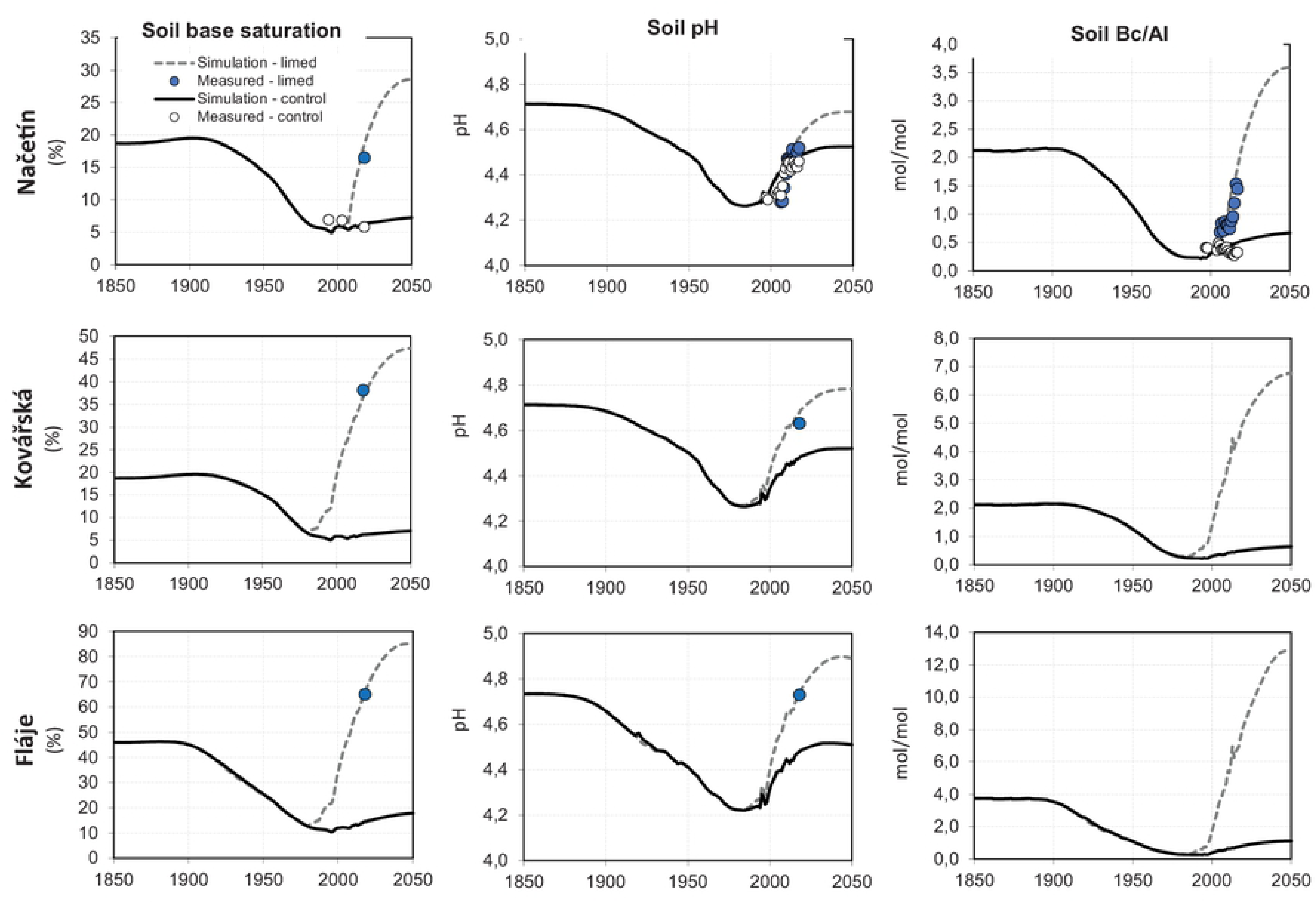
Modelled and measured (a) soil base saturation, (b) soil solution pH and (c) soil solution Bc/Al ratio (1850-2050) for all investigated sites.

Preindustrial soil water pH was estimated to be 4.7 (Fig. 8) at Načetín, and the lowest pH (4.25) was modelled for the 1980s. pH started to rise soon after the decline in deposition in the 1990s (Fig. 5). A pH of 4.45 was reached at the naturally regenerated control plot by 2018 (Table II). The limed plot’s pH increased to 4.55 between 2007 and 2018. Modelled was 4.7 for the limed plot and 4.5 for the control plot. Almost identical preindustrial (4.7) and minimum pH (4.25) pH values were estimated for Kovářská. Liming since 1981 resulted in measured and modelled pH of 4.65 in 2018 and an estimated pH of 4.8 in 2050. Highly weathered basaltic and limed Fláje showed very similar patterns as Kovářská (Fig. 8.) – preindustrial pH=4.75, minimum 4.25 and present pH=4.7. The estimated pH for 2050 (4.9) was higher than the preindustrial estimate.

The molar Bc/Al ratio ((Ca + Mg + K)/Al) in soil solution was estimated to be around two as a preindustrial value (Fig. 8) and it declined to as low as 0.2 in the 1980s. In contrast to pH, it increased only slightly and stay around 0.4 between 1994-2018. At the limed plot, Bc/Al increased significantly to 1.5 in 2018 and future predictions are that Bc/Al will rise to 3.5 in 2050. A similar pattern was modelled for Kovářská, where Bc/Al was estimated at 4.5 in 2018 (from a minimum of 0.2 in the 1980s before liming) and almost 7.0 was predicted for 2050 after liming. At base-rich and limed Fláje site, preindustrial Bc/Al was estimated at 3.9, with a minimum of 0.2 for the beginning of the 1980s, 8.0 for 2018 and 13 for 2050.

### 4.3. TRW chronologies

The site TRW chronologies from the four sites vary in length from 59 to 106 tree rings. The high reliability of all site chronologies was confirmed by the Rbar (>0.48) and EPS (>0.92) values, which remained above the threshold of 0.85 [39] for the entire study period. The high similarity among TRW chronologies at each site allows compilation of the mean TRW chronology (Rbar=0.40, EPS=0.97), covering the period from 1912 to 2017 (Fig. 9A).

**Figure 9.**
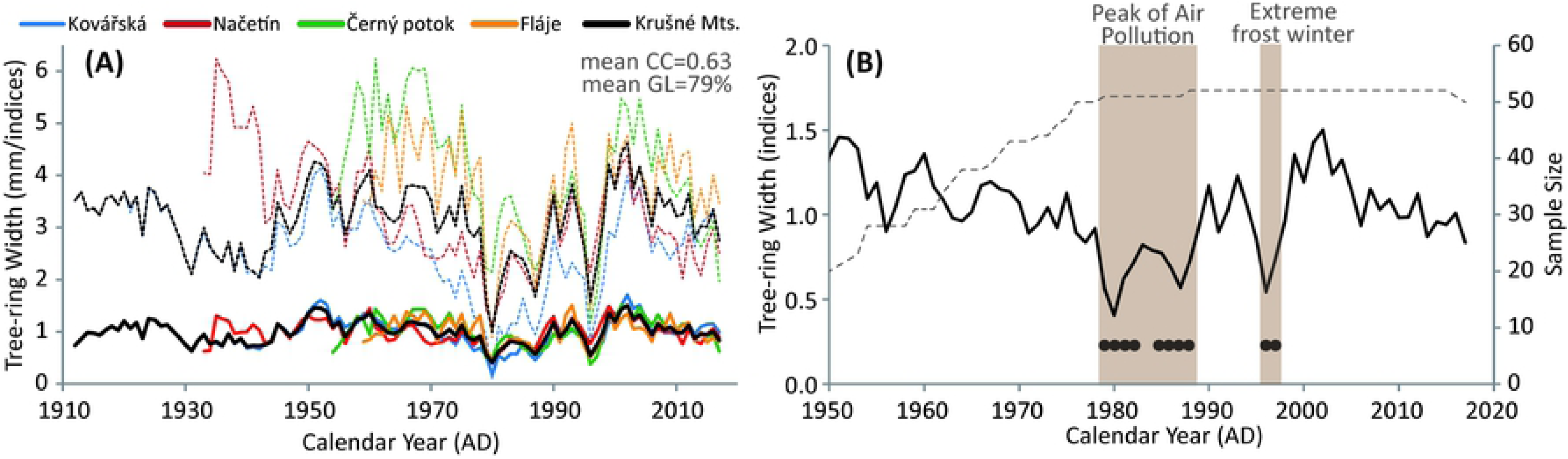
(A) Raw (dotted line) and standardised (full line) TRW chronologies of all individual sites as well as Ore (Krušné) Mts. Mean correlation coefficient (CC) and mean GL (Gleichläufgkeit; Eckstein, Bauch 1969) indicate similarity among the site chronologies. (B) Indexed mean TRW chronology for Ore Mts. truncated for sample size (dotted line) of at least 20 TRW series. Black dots indicated negative extremes.

Replication of the mean chronology decreased backwards and dropped below the 20 TRW series before 1950 (Fig. 9B). The mean chronology with an average annual growth rate of 2.75 mm shows a considerable decrease in annual growth starting in the 1950s. Three most significant growth reductions were revealed by the analysis of negative extremes in the periods 1979–1982, 1985–1988, and 1996–1997 when average tree-ring widths drop to 1.42, 1.67, and 1.49 mm, respectively. Additionally, we detect seven missing rings between 1979–1982 and five in 1996–1997. The interval of the most pronounced growth reduction (1979–1988), likely initiated by the extremely cold and harsh winter of 1978/1979 [40], corresponds also to one of the highest concentrations of SO_2_ (Fig. 3).

Radial growth of conifers at high altitudes is primarily driven by growing season temperature and global radiation [41]. However, the strong growth–temperature relationship was significantly reduced in the second half of the 20^th^ century when mountain forests of Central Europe experienced widespread and long-lasting effects related to acid deposition [42]. The coincidence of freezing temperatures in winter 1978/1979 and high concentrations of SO_2_ led to tree-ring width fluctuations and the weakening of the climate signal. In the 1980s, the radial growth of Norway spruce in the “Black Triangle” was not controlled by summer temperature at all. Extremely narrow tree-ring widths or even missing rings were detected under exceptionally unfavourable environmental conditions [14].

During the time of the highest sulfur emissions, the TRW reached its lowest values (Fig. 9). The TRW indices were reduced by >40% during the 1980s compared with their long-term mean (1950–2018). After the decline in sulfur deposition, the mean TRW indices increased, exceeding shortly the values from the period before the extreme emission load.

### 4.4. Relationships between forest growth and acidification

Direct (SO_2_ concentrations, atmospheric deposition) and indirect (soil chemistry represented by soil pH, Bc/Al and soil base saturation) factors were evaluated to judge their relevance in the retrospectively observed reduced TRW (Fig. 9.).

#### 4.4.1. Atmospheric deposition and SO_2_ concentrations

Atmospheric deposition of sulfur was identified as the most critical factor controlling reduced TRW for the period of tree lifetimes (Table III) for gneiss underlain Načetín (since 1933) and Kovářská (since 1912). S deposition was not identified to be important for reduced growth at basalt underlain Fláje (since 1959). S deposition was the only relevant explanatory parameter when the whole tree lifetime was analysed. TRW was reduced since 1979 (Fig. 9) and has been fully recovered since 2000. It represented deposition of about 300-350 meq·m^-2^·yr^-1^ at the beginning of reduction and ca. 250 at the end of the TRW reduction period (S1 Annex, Fig. S1.3.1).

**Table III.** Correlation matrix (R) and regression coefficients (R^2^) between the individual parameters and TRW. Two intervals were correlated: (i) modelled annual atmospheric deposition and soil chemistry for the life span of trees and (ii) modelled annual soil chemistry, deposition and measured annual airborne SO_2_ concentrations for the period 1979-2017. Statistically significant (p<0.05) values are displayed in red bold.

|  | 1850-2017 |  | 1979-2017 |  |
| --- | --- | --- | --- | --- |
| **Načetín** | R | R<sup>2</sup> | R | R<sup>2</sup> |
| Total S deposition | **-0.40** | **0.16** | **-0.49** | **0.24** |
| Air SO<sub>2</sub> imissions |  |  | **-0.50** | **0.25** |
| Soil pH | 0.23 | 0.05 | 0.18 | 0.03 |
| Soil base saturation | 0.13 | 0.02 | -0.30 | 0.09 |
| Soil Bc/Al | 0.14 | 0.02 | 0.06 | 0.00 |
| **Kovářská** |  |  |  |  |
| Total S deposition | **-0.38** | **0.14** | **-0.72** | **0.51** |
| Air SO<sub>2</sub> imissions |  |  | **-0.69** | **0.48** |
| Soil pH | 0.28 | 0.08 | **0.56** | **0.31** |
| Soil base saturation | 0.34 | 0.11 | **0.59** | **0.35** |
| Soil Bc/Al | 0.26 | 0.07 | **0.49** | **0.24** |
| **Fláje** |  |  |  |  |
| Total S deposition | -0.14 | 0.02 | **-0.45** | **0.20** |
| Air SO<sub>2</sub> imissions |  |  | **-0.45** | **0.20** |
| Soil pH | 0.07 | 0.01 | 0.30 | 0.09 |
| Soil base saturation | 0.07 | 0.00 | 0.34 | 0.11 |
| Soil Bc/Al | 0.02 | 0.00 | 0.23 | 0.05 |

TRW was additionally and deeply reduced in the years 1996-1997. Acid rime with extremely low pH was recorded during winter 1995/1996 [36]. It resulted in significant defoliation in spring 1996 and a subsequent decrease of TRW for two years. Trees fully recovered their radial growth in 1998. The acid rime episode was caused by an asymmetrical reduction of dust and SO_2_ emissions (Fig. 2). Dust from power plants was reduced earlier and more effectively than SO_2_ (Fig. 2). Thus precipitation and rime were extremely acidic in this short period in the mid-1990s. TRW reduction was observed at all sites (Fig. 9). Still, the most pronounced reduction was observed at the oldest stand at Kovářská and the lowest reduction was observed at the youngest stand, Fláje, underpinning the role of the vitality associated with young stands.

SO_2_ concentrations in the air were available for the period 1979-2017 from the Zinwald station (German-Czech border). This record was the longest measured at one station in the region. Clearly, it represented the level of air pollution in the “Black triangle” region where Czech, as well as German and Polish power plants, contributed to the extreme local pollution [2]. Analysing relationships for 1979-2017 only, SO_2_ concentrations and sulfur deposition explained forest decline equally and statistically significant at all sites (p<0.05, Table III.). Both auto-correlated values (as S-SO_4_ deposition is derived from the oxidation of SO_2_ in the atmosphere) explain TRW better at naturally acidic plots with oldest spruce stands (Načetín, Kovářská) followed by alkaline Fláje with a young spruce stand. The possible explanations are that (i) young stands are more vital (only 30 years in 1979 compare to ca. 55 at Načetín and 80 years at Kovářská), or (ii) base-rich bedrock better supplied spruce needles with nutrient cations (Ca, Mg, K) and as a result, they were more resistant to their leaching by acid rain and subsequent needle injury and defoliation [12].

#### 4.4.2 Soil chemistry

Soil chemistry did not show a statistically significant correlation with TRW for the lifetime of trees at any site (Table III.). It also did not explain changes in tree growth changes shorter periods with rapid changes in pollution (1979-2017) for Načetín control and Fláje. The only site where soil chemistry, as well as air pollution, explains TRW is Kovářská. This site is naturally acidic (gneiss in bedrock), and it was first time limed in 1981 (Table I), during the most profound TRW depression (Fig. 9B). Thus soil solution pH, Bc/Al and base saturation increased significantly and monotonically (Fig. 8) from low values (BS of 6% in 40 cm) to very high saturation (38% in 2018), significantly higher than the preindustrial estimate of 20%. Such artificial soil treatment makes soil chemistry “recovery” very robust, linear and coincident with a declining level of air pollution. At acidic and unlined Načetín, as well as at naturally alkaline and limed Fláje, TRW did not correlate with modelled soil chemistry.

## 5. Synthesis

Long-term changes in soil chemistry (soil pH, base saturation and Bc/Al ratio) did not explain the observed decline in TRW at both of our sites for which data are available (Table III) despite the depletion of nutrient cations and enhanced concentration of potentially toxic Al in the soil solution that has frequently been hypothesised to lead to deterioration of forest health. The molar Bc/Al ratio ((Ca + Mg + K)/Al) has been widely used as a criterion for the risk of tree damage [43, 44, 45]. Experiments with seedlings by Sverdrup et al. [43] showed that increased mortality occurred if the Bc/Al ratio was lower than 1. Field data from the Czech Republic [45] suggested that increasing tree damage occurred with decreasing Bc/Al in the soil solution of the rooting zone in Norway spruce stands. On the other hand, this concept’s limitation was shown by De Witt et al. [46]. They found that of the base cations, only reduction of Mg uptake occurred after long-term experimental addition of AlCl_3_ to the rooting zone of Norway spruce in southern Norway. Our observations support the hypothesis that direct injury of the assimilatory organs is more important for tree damage, at least in the areas where extremely high SO_2_ concentrations, as well as S deposition, occurred in the past [e. g. 1, 14, 42].

If the shorter period 1979-2017 is examined (Table III), statistically significant correlations for soil pH, base saturation, and Bc/Al were observed for Kovářská only (Table III). This plot was limed three times since 1981 (Fig. 8), and soil chemistry became less acidic and base-rich than the unlimed Načetín control (the same bedrock), where no correlation was observed. The magnitude of soil chemistry change at Kovářská was very pronounced in comparison to Načetín control (Fig. 8), but TRW changes were similar and no statistical difference between TRW recovery after 1979 was detected (Fig. 9).

The minor role of soil chemistry was also manifested at naturally well buffered and also massively limed Fláje. Very high base saturation before the liming and supersaturation after that (Fig. 8) did not eliminate TRW decline in the 1970s and 1980s (Fig. 9).

Similar findings from Krkonoše Mts. National Park (ca. 150 km east of our sites) were published by Kolář et al. [14]. They found that TRW of Norway spruce declined significantly at high altitudes since the 1970s and recovered fully around 2000 when acidic deposition declined. One plot from five investigated was limed by 5 t·ha^-1^ of dolomitic limestone in the mid-1980s. Liming did not affect TRW, and all five sites recovered synchronously as the acid deposition declined.

The fact that canopy injury was the most important for TRW reduction was well documented during the frost/acid rime episode (Fig. 4.) in the winter of 1995/1996 (Fig. 9). It was followed by a short but deep depression in the tree ring width, even though soil chemistry did not show any deviation from the observed long-term trends (Fig. 6 and Fig. 8).

Our results suggest that the direct impact of acidic deposition from high SO_2_ concentrations and/or sulfur deposition was the main driver of forest decline. Soil chemistry to a depth of 40 cm most probably did not play a crucial role in the observed forest dieback in the heavily polluted so-called “Black Triangle” of Central Europe, formerly one of the most SO_2_-polluted areas in the world. Soil liming did not appear to help recover tree growth from chronic stress as has been suggested and often used as an argument for repeated liming [37]. Liming may help reduce soil acidity and increase the amount of deficient base cations (Ca and Mg). Still, concerning tree growth measured as changes in TRW, our study showed no positive effects of liming. Other authors [e. g. 47] have also found that enhanced radial growth of trees did not occur after dolomitic limestone application.

We cannot reject altogether the hypothesis that forest growth might be negatively affected by soil chemistry, as the uppermost organic and mineral soil down to 10 cm slightly recovered at long-term observed Načetín control plot (Fig. 6). In contrast, deeper mineral soil down to 40 cm became even more acidic during the last few decades (Fig. 6). Very acidic forest soils (down to 30 cm) were detected at 31% of 1599 semi-randomly selected soil samples taken between 2006-2009 in the Czech Republic [48], but significant forest decline which can be attributed to soil acidification was not observed during the same time [49]. As roots of Norway spruce are located mainly in the organic layer and upper mineral soil, more favourable conditions (less Al, slightly higher soil base saturation) were created in the 1990s. But the uppermost layers responded quickly to the steep decline of acidic deposition (Fig. 5). Thus it is difficult to disentangle the effects of atmospheric chemistry and rapid but limited soil recovery in the uppermost rooting zone.

As has been shown in this paper and many papers and reviews previously [e. g. 38, 50, 51], acidic deposition caused a deterioration in soil chemistry, but negative effects on Norway spruce growth did not occur. Direct effects on foliage, reducing primary productivity, appear to be more critical for reducing tree growth than soil acidification.

## Acknowledgements

This work was financially supported by Czech Science Foundation (GAČR) grant 18-17295S and *SustES – Adaptation strategies for sustainable ecosystem services and food security under adverse environmental conditions (CZ.02.1.01/0.0/0.0/16_019/0000797)* granted by Czech Ministry of Education, Youth and Sport. Pavla Holečková from Czech Geological Survey is thanked for the editorial assistence.

---

**S1 Annex FigureS1.1.1.**
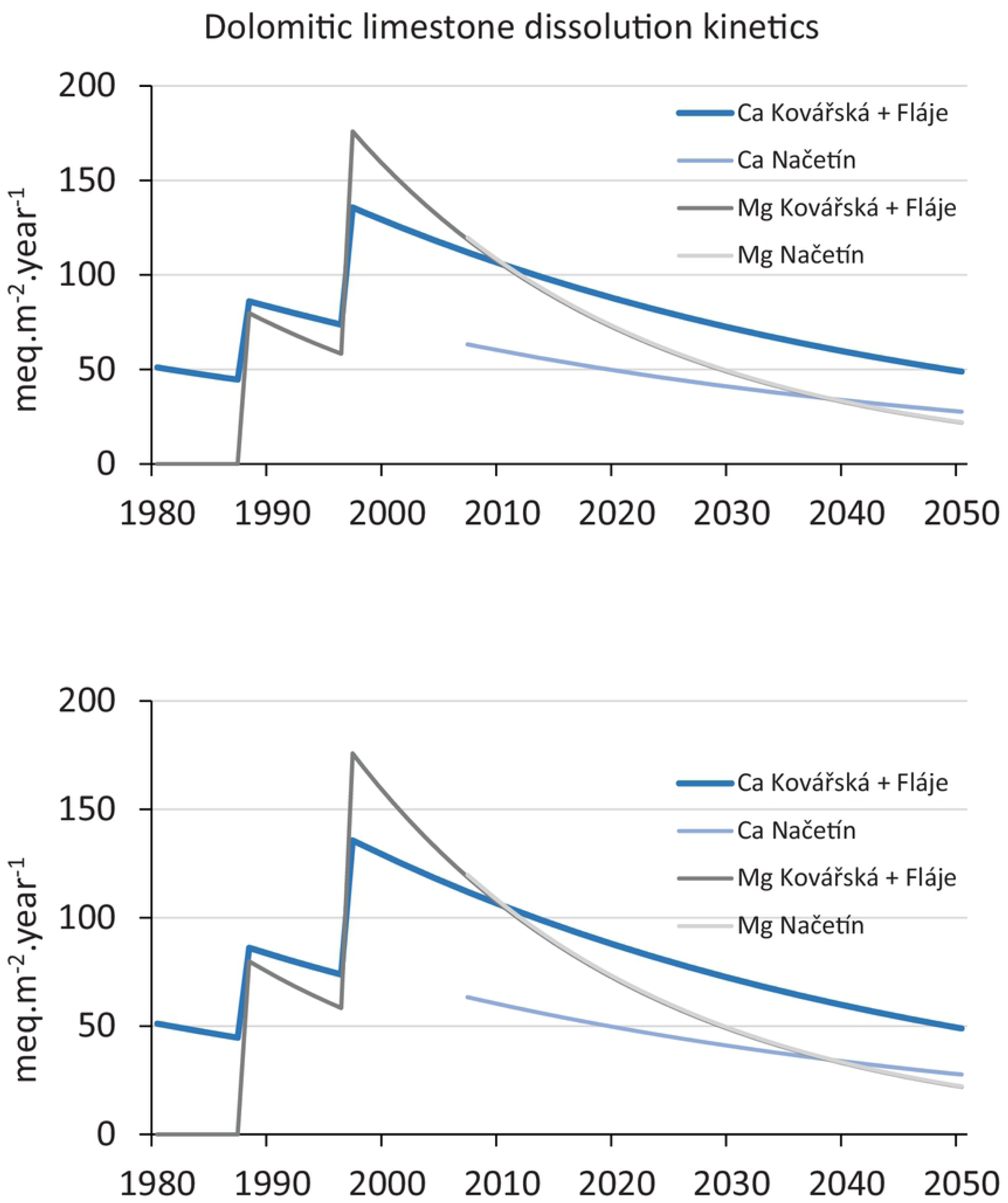

**S1 Annex FigureS1.2.1.**
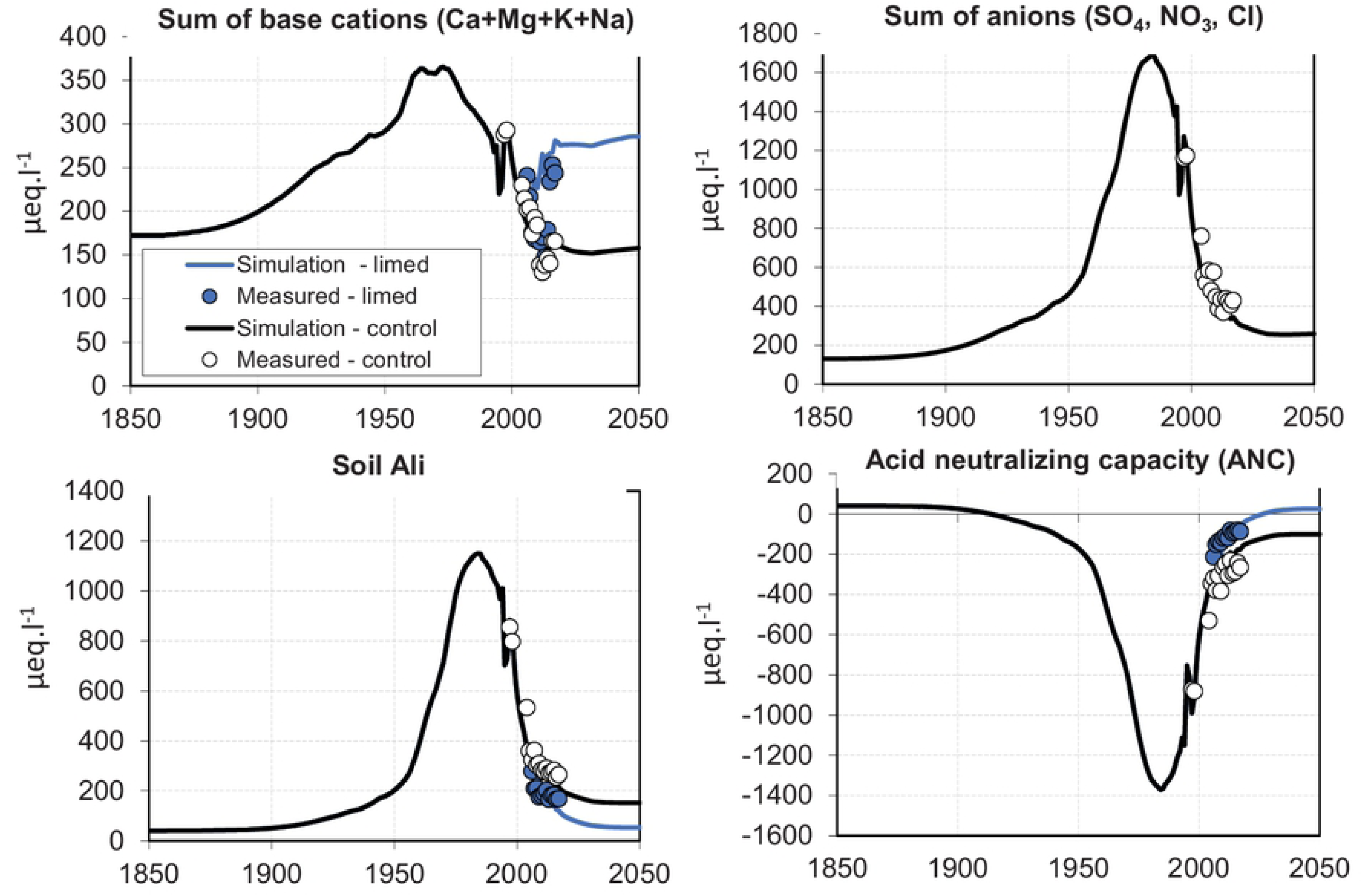

**S1 Annex FigureS1.2.2.**
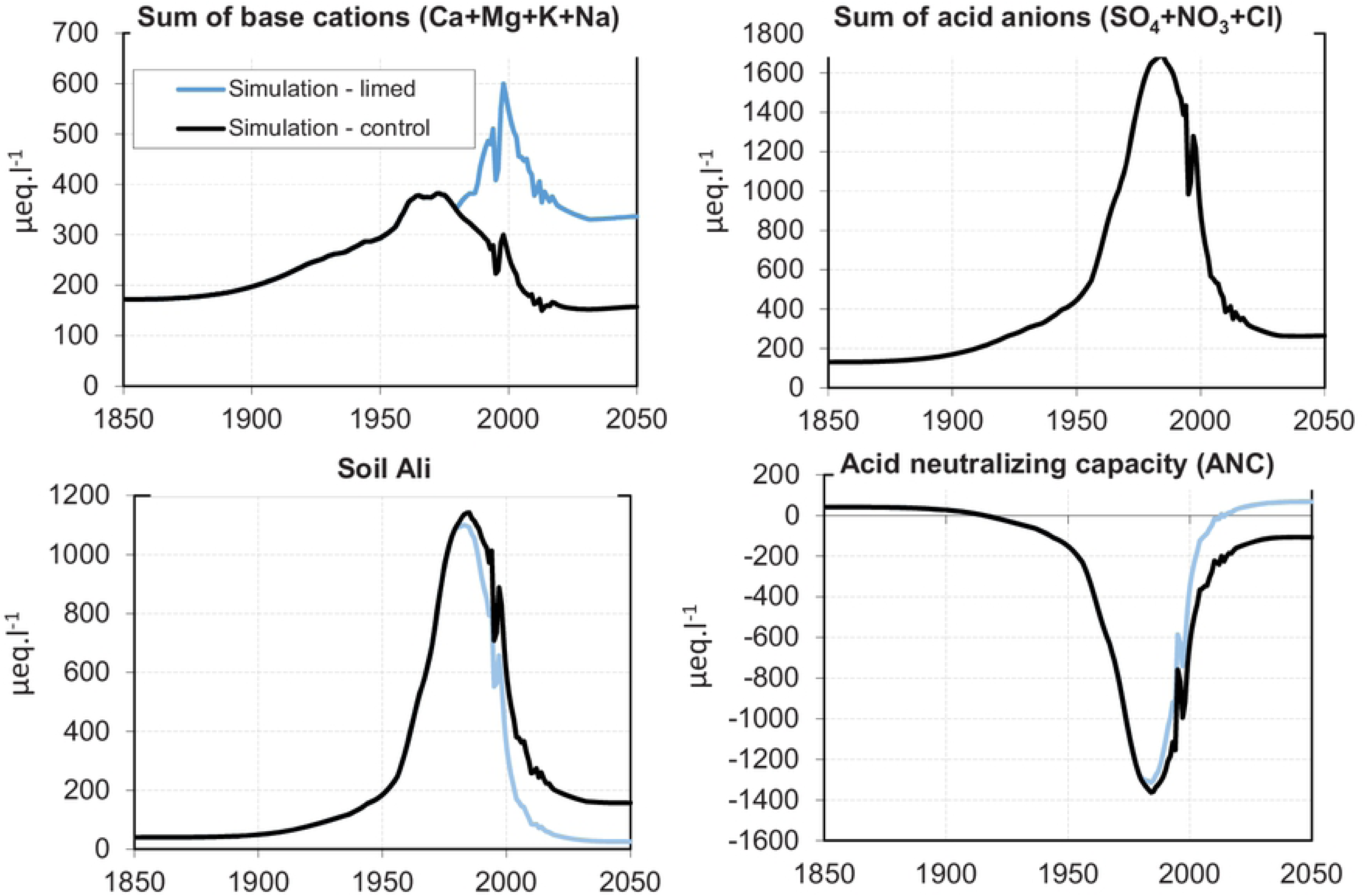

**S1 Annex FigureS1.2.3.**
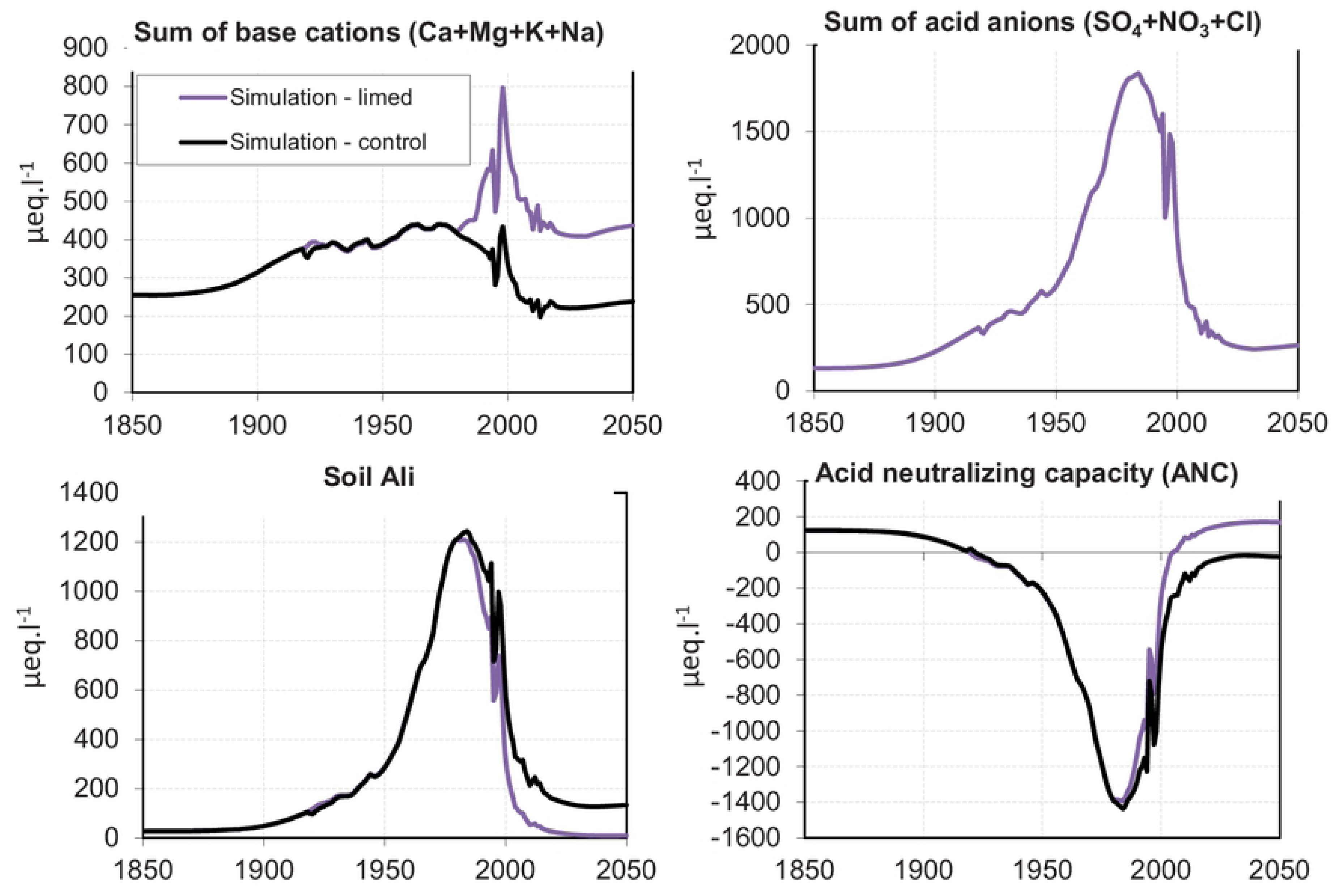

**S1 Annex FigureS1.3.1.**
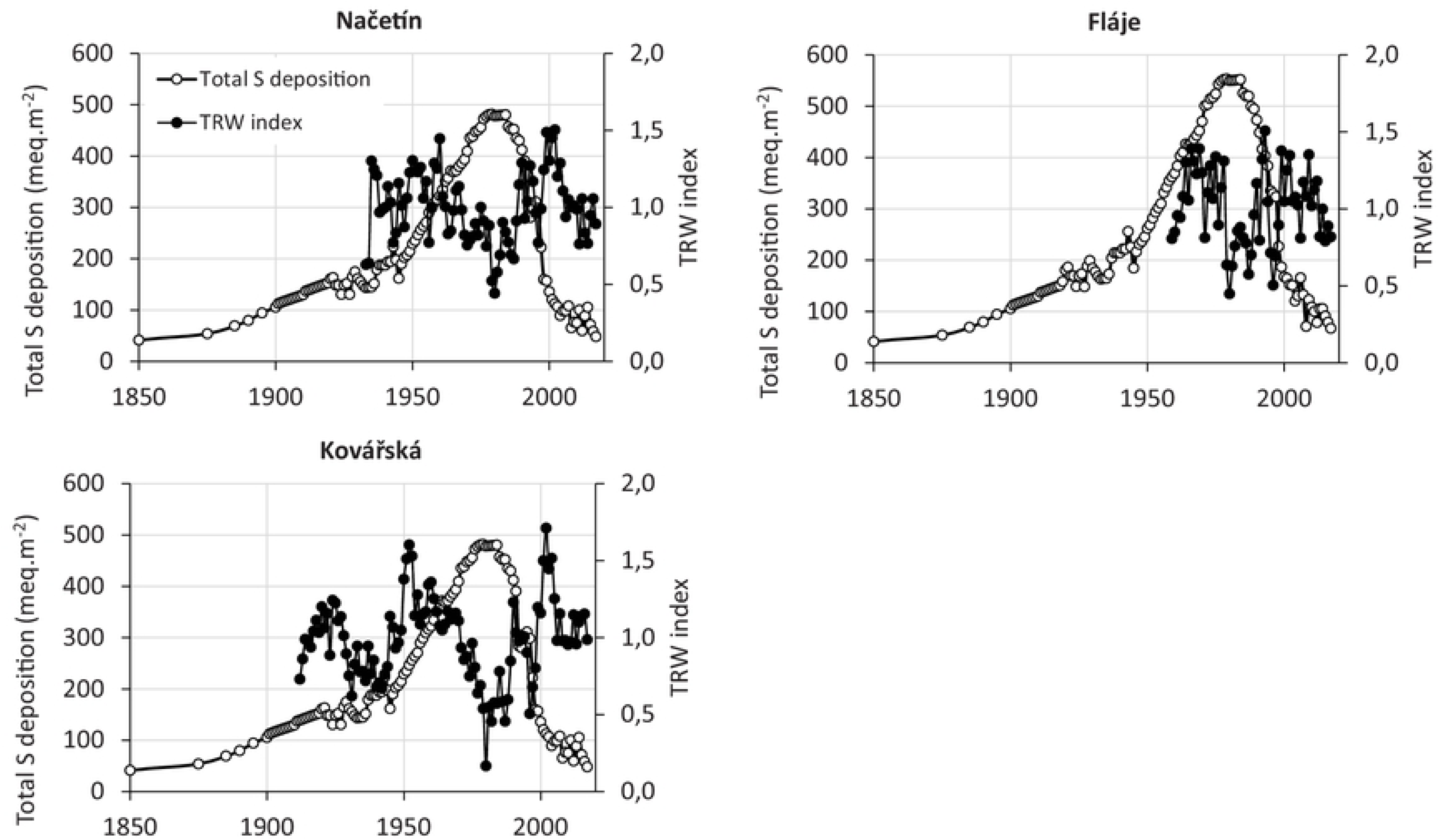

